# Expected reward value and reward prediction errors reinforce but also interfere with human time perception

**DOI:** 10.1101/2024.04.17.589985

**Authors:** Emily K. DiMarco, Ashley Ratcliffe Shipp, Kenneth T. Kishida

## Abstract

Time perception is often investigated in animal models and in humans using instrumental paradigms where reinforcement learning (RL) and associated dopaminergic processes have modulatory effects. For example, interval timing, which includes the judgment of relatively short intervals of time (ranging from milliseconds to minutes), has been shown to be modulated by manipulations of striatal dopamine. The ‘expected value of reward’ (EV) and ‘reward prediction errors’ (RPEs) are key variables described in RL-theory that explain dopaminergic signals during reward processing during instrumental learning. Notably, the underlying connection between RL-processes and time perception in humans is relatively underexplored. Herein, we investigated the impact of EV and RPEs on interval timing in humans. We tested the hypotheses that EV and RPEs modulate the experience of short time intervals. We demonstrate that expectations of monetary gains or losses increases the initial performance error for 1000ms intervals. Temporal learning over repeated trials is observed with accelerated learning of non-reinforced 1000ms intervals; however, RPEs – specifically about rewards and not punishments – appear to reinforce performance errors, which effectively interferes with the rate at which (reinforced) 1000ms intervals were learned. These effects were not significant for 3000ms and 5000ms intervals. Our results demonstrate that EV and RPEs influence human behavior about 1000ms time intervals. We discuss our results considering model-free ‘temporal difference RL-theory’, which suggests the hypothesis that interval timing may be mediated by dopaminergic signals that reinforce the learning and prediction of dynamic state-transitions which could be encoded without an explicit reference to ‘time’ intervals.

## INTRODUCTION

The neurobiological mechanisms underlying the human experience of time and the execution of timing dependent behaviors remains elusive (Buhusi & Meck, 2005; Patton & Buonomano, 2018). Considerable evidence suggests that for relatively short time intervals, dopaminergic signals are critical (Meck, 1996; Petter, et al., 2018). To date, the best account of what dopamine neurons encode – and what dopamine release signals – lies with computational reinforcement learning theory (Sutton & Barto, 1998; Montague, et al., 1996; Shultz, et al., 1997; Uchida, et al., 2022; Sands, et al., 2024) within the particular theoretical framework, ‘temporal difference reinforcement learning’ (TDRL), explaining dynamic changes in dopamine neuron activity and changes in dopamine release throughout the course of learning Pavlovian and Instrumental associations (Montague, et al., 1996; Shultz, et al., 1997; Kim, et al., 2020; Sands, et al., 2023). An explanation of dopamine’s role in timing behavior ought to be possible in the context of TDRL’s theoretical terms; however, little work has been done to explicitly investigate TDRL defined variables in the context of timing behavior (though see Gershman, et al., 2014; Petter, et al., 2018; Mikhael & Gershman, 2019; and Jakob, et al., 2022). Here we investigated interval timing behavior and temporal learning while varying reward and punishment reinforcement and analyzed our data with a particular aim to disentangle TDRL terms and the corresponding hypothesized role for dopamine.

Many reports have linked dopaminergic processes with timing behavior about relatively short intervals of time - on the order of hundreds of milliseconds to several seconds (Buhusi & Meck, 2005). These behaviors are typically modelled in humans and rodents using interval timing tasks like peak interval procedures (**Fig 1**, Rakitin et al., 1998; Balci & Freestone, 2020). In humans, the task can be verbally instructed with little dependence on *de novo* learning required, though improvement in performance (i.e., temporal learning) can be observed over repeated trials. In rodents, however, as well as any other non-human animal model, interval timing behavior must first be trained using instrumental reinforcement learning methods that utilize rewards to reinforce desired behaviors. Work attempting to use animal models to disentangle the role dopamine plays in the mechanisms underlying laboratory interval timing behavior face the confound that dopaminergic processes and reward-based learning is required to train and elicit timing behaviors. Humans, on the other hand, can be instructed and are able to perform tasks without motivational drive provided by extrinsic rewards, which allows the comparison of reinforced versus non-reinforced interval timing behaviors and the impact that learning signals have on temporal learning and interval timing behavior.

**Fig 1.**
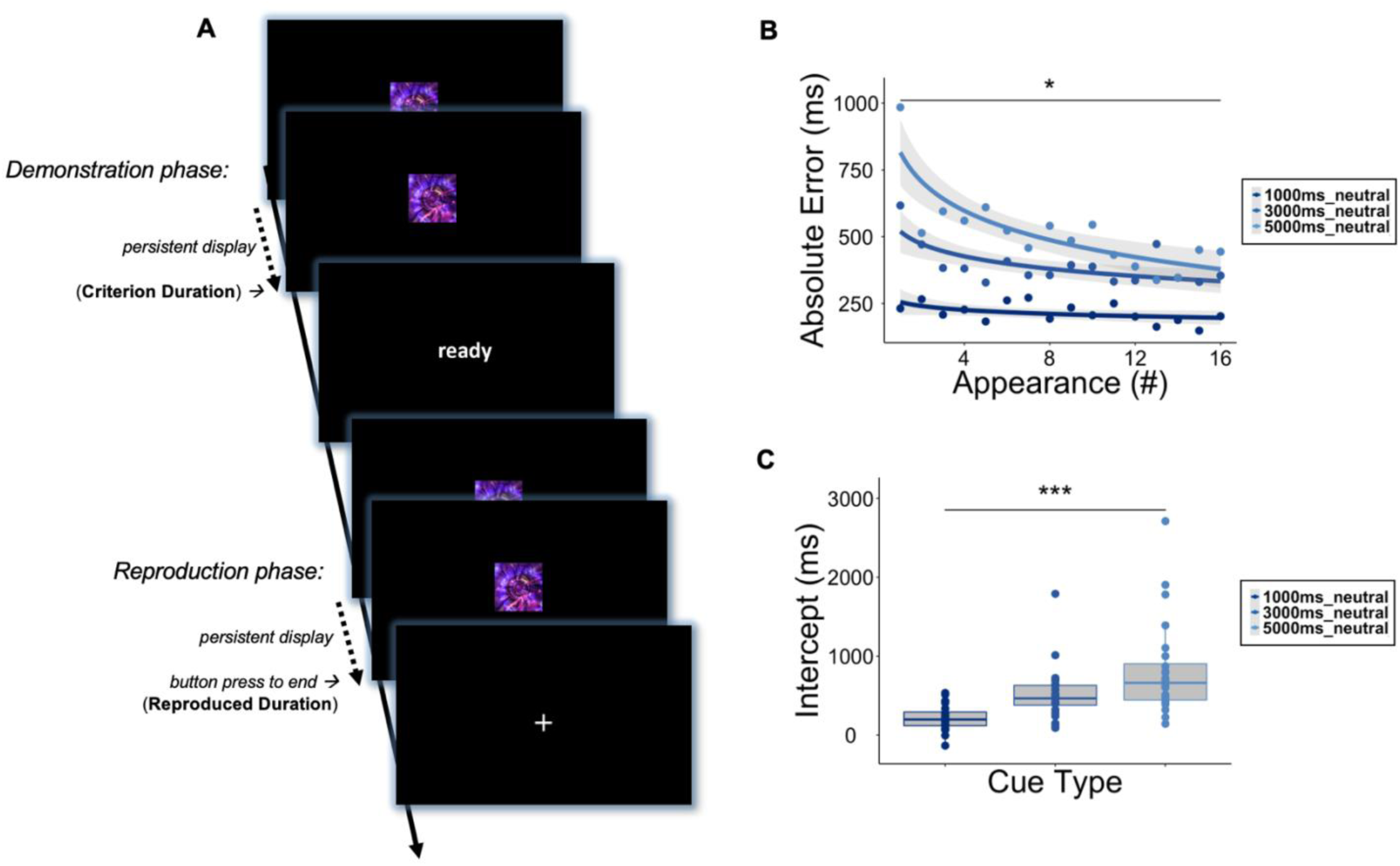
Significant temporal learning occurs in response to non-reinforced cues. **A.** A trial of the PIP-n (PIP-neutral) began with a ’Cue Onset’, where a cue was presented on the screen for a criterion duration (CD; 5000ms, 3000ms, or 1000ms), indicated by distinct cues. Following the presentation of a CD, a black screen appeared for 500ms followed by a ‘ready’ prompt (fixed 1000ms) to prepare the participant to reproduce the cued CD. Following another 500ms black screen, the cue was then re-presented and remained on the screen until the participant reproduced the cued CD with a button press, or a time twice the CD passed. **B.** Learning curves were calculated for each participant (n = 24) based on absolute error for each cue type at each appearance. Mean absolute error at each appearance was then fit with an exponential function to determine the rate of temporal learning overtime. Learning curves for the representative neutral-2 cues on the PIP-n show significantly lower error at appearance 16 when compared to appearance one. **C.** When comparing intercepts of learning curves across criterion durations, intercepts increased as CDs increased, revealing increasing error for larger intervals. *Significance based on p < 0.05*, p < 0.01**, p < 0.001***. Shading = SEM*.

Through a variety of positive or negative reinforcement schedules, a wide range of Pavlovian and Instrumental learning effects can be trained and investigated in animal models and humans (MacInnis & Guilhardi, 2006). Amongst the most widely utilized approaches are those that use positive reinforcement via delivery of rewards to reinforce certain behaviors or environmental states. Both passive (i.e., Pavlovian) and active (i.e., Instrumental) learning have been shown to involve dopamine signaling; and, under both psychological constructs, time intervals were shown to be learned with accuracy and precision (Church, 2012). It has also been shown that dopamine neurons in a non-human primate model respond in a manner consistent with a theorized optimal teaching signal called a “temporal difference reward prediction error” during Pavlovian conditioning (Montague, et al., 1996; Shultz, et al., 1997). Notably, dopamine neuron responses show precise time encoding as evidenced by the ‘pause’ in firing activity (indicating a negative reward prediction error) when – after learning – a reward was expected at a particular time but not delivered (Montague, et al., 1996; Shultz, et al., 1997). This result, suggesting that dopamine neurons and dopamine release encode a “temporal difference reward prediction error” during instrumental learning behavior has been replicated a number of times (Montague, et al., 2004; Glimcher, 2011; Kishida & Sands, 2021) in rodents (Uchida, et al., 2022) and in humans (Sands, et al., 2023).

Dopamine’s role in timing behavior has been implicated by several complementary approaches in animal models and in humans. Neuropharmacological methods in human studies have shown that dopamine depletion (Coull et al., 2012), dopamine receptor antagonists (Drew et al., 2003; Buhusi & Meck, 2002) and psychostimulants that affect the dopamine system (Buhusi & Meck, 2002) alter timing behavior and time perception(Coull et al., 2012; Drew et al., 2003; Buhusi & Meck, 2002; Rammsayer, 1999). Wiener and colleagues identified an association between dopaminergic system genes and timing in humans (Wiener, et al., 2011). More recently, it has been demonstrated that rapid changes in dopamine neuron activity in mice (Jakob et al., 2022), manipulation of dopamine neuron activity using optogenetic methods in mice (Soares et al., 2016), and rapid (i.e., sub-second) and slow (i.e., several seconds) changes in dopamine levels in humans affects timing behavior and time perception (Sadibolova, et al., 2024; Sadibolova, et al., 2022; Terhune, et al., 2016). Notably, many of the methods used to implicate, measure, or manipulate dopamine levels during instrumental timing behavior is also used to investigate dopamine’s role in instrumental (and Pavlovian) reinforcement and learning (Frank & Fossella, 2011; Kishida & Sands, 2021). This has led us and others (Petter, et al., 2018; Mikhael & Gershman, 2019) to hypothesize that dopaminergic timing behavior may be mediated by the same mechanisms that underlie dopaminergic reinforcement learning.

In this study, we explored the role reinforcement learning signals (i.e., ‘expected value of reward’ (EV) and reward prediction errors (RPEs)) play in interval timing behavior in humans. Recently, these behavioral signals were shown to elicit a phasic (sub-second) change in dopamine levels in the human striatum during an instrumental learning task that did not modulate time intervals but did possess an explicit temporal structure (Sands, et al., 2023). We note that, according to the temporal difference RL framework, cues that predict a reward also elicit an EV-driven RPE, which is not described by the solely extrinsic-reward driven RPE described Rescorla-Wagner framework (Rescorla & Wagner, 1972). Thus, we hypothesized that changes in the EV of ‘criterion duration’ cues, associated RPEs, and RPEs following execution of timing behaviors with extrinsic reward or punishment would each modulate interval timing behavior in a manner consistent with expected changes in striatal dopamine levels.

To test our hypotheses, we executed two versions of a peak interval procedure (PIP) that investigated the reproduction of 1000ms, 3000ms, and 5000ms intervals of time (**Fig 1A**, **2A**). In the first task (**Fig 1A**), 24 participants (**Table 1**) completed ‘PIP-neutral’ (PIP-n), which tested the reproduction of demonstrated criterion durations in the absence of any reinforcement. Next, 24 naïve participants (**Table 1**) completed ‘PIP-reinforced’ (PIP-r, **Fig 2A**), which provided positive, negative, or no monetary reinforcement that scaled linearly with the accuracy of reproduction of the criterion durations (**Fig 2B**). We compared timing behavior in the PIP-neutral task to timing behavior in the PIP-reinforced task. Specifically, we determined and compared the rate of temporal learning when extrinsic (i.e., monetary) reinforcement was provided (PIP-r) compared to when it was absent (PIP-n). We also assessed the impact of RPEs on the subsequent performance of the same criterion durations in the subsequent trial. We report significant effects of the EV on accelerating temporal learning (**Fig 3**). However, we also demonstrate that reinforcement can paradoxically interfere with temporal learning (**Fig 4**). We discuss our results considering model-free ‘temporal difference RL-theory’ and discuss how our results and this theoretical framework suggest the hypothesis that the brain can use dopaminergic reinforcement to learn ‘time’ intervals by simply encoding dynamic state-transitions, which, notably, would not require a ‘pacemaker’ or ‘internal clock’ to keep time.

**Fig 2.**
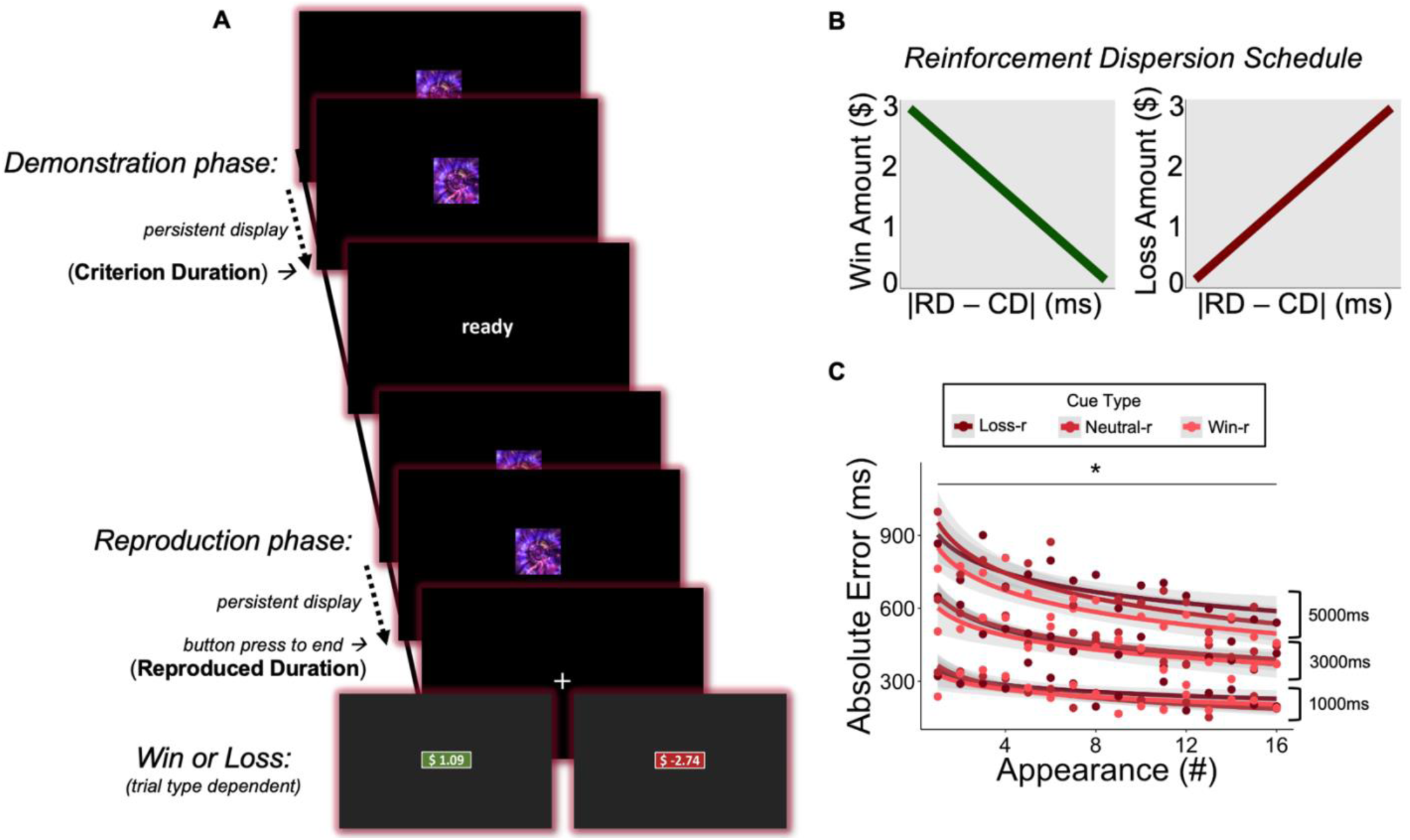
Significant temporal learning occurs in response to reinforced cues. **A.** A trial of the PIP-r (PIP-reinforced) began with a ’Cue Onset’, where a cue was presented on the screen for a criterion duration (CD; 5000ms, 3000ms, or 1000ms), indicated by distinct cues. Following the presentation of a CD, a black screen appeared for 500ms followed by a ‘ready’ prompt (fixed 1000ms) to prepare the participant to reproduce the cued CD. Following another 500ms black screen, the cue was then re-presented and remained on the screen until the participant reproduced the cued CD with a button press, or a time twice the CD passed. Reinforcements (win or loss) were presented following the button press (fixed 500ms). **B.** Reinforcements were linearly scaled to the absolute difference between the reproduced duration (RD) and the CD, with a maximum possible reward or loss of ±$3 **C.** Learning curves were calculated for each participant (n = 24) based on absolute error for each cue type at each appearance. Mean absolute error at each appearance was then fit with an exponential function to determine the rate of temporal learning overtime. Learning curves for each cue type on PIP-r show significantly lower error at the last appearance (16) when compared to the first appearance (1), for all cues. *Significance based on p < 0.05*. Shading = SEM*.

**Fig 3.**
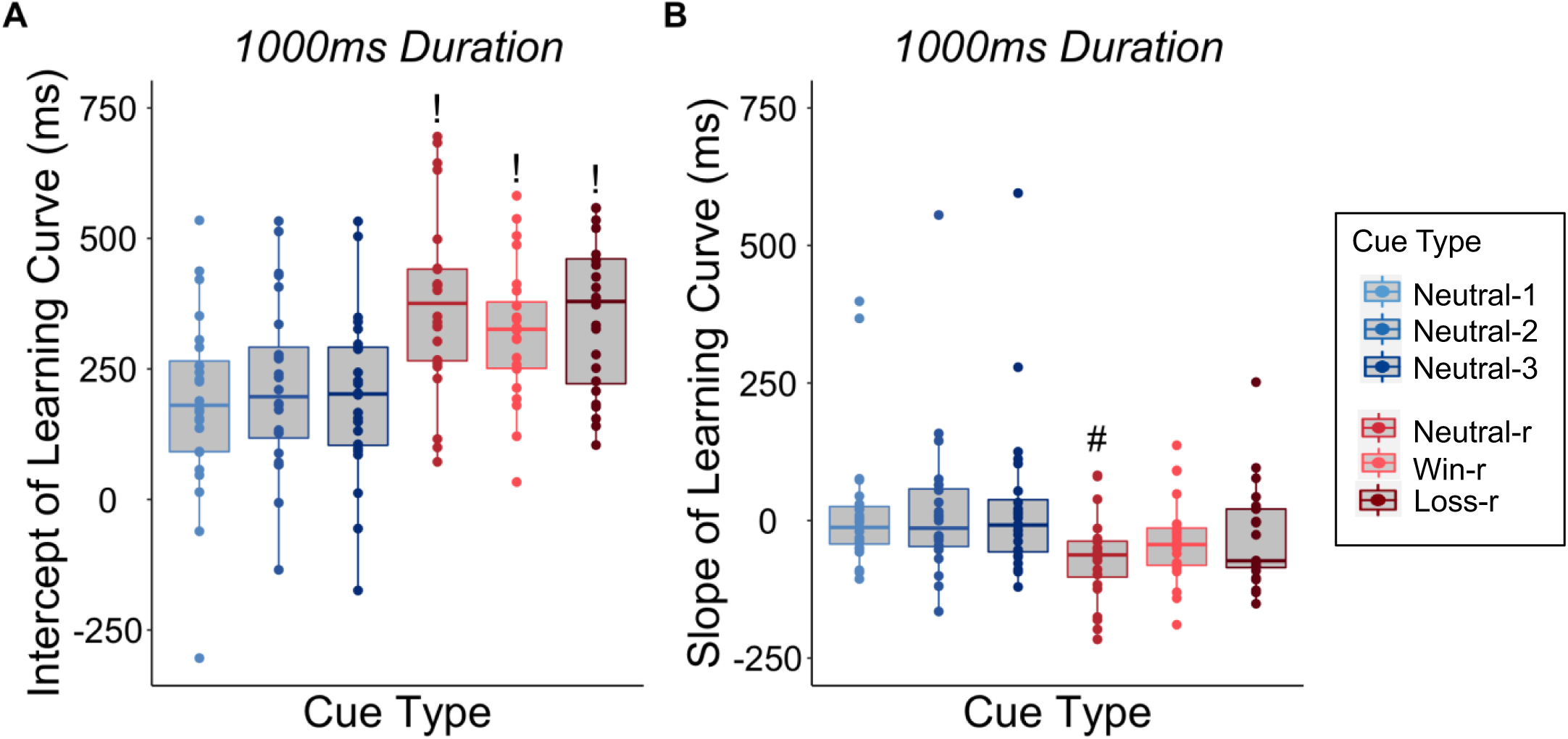
Reinforcements increase the intercepts of the learning curves, which generalizes to the neutral cues on 1000ms duration. **A.** The intercepts of the learning curves are significantly different between the neutral cues (Neutral-1, Neutral-2, Neutral-3) on the PIP-n and all cues (Neutral-r, Win-r, Loss-r) on the PIP-r for the 1000ms criterion duration. *“!” denotes group is significantly different than all three neutral groups (blue) based on pair-wise t-tests.* **B.** The slopes of the learning curves are significantly different between the neutral cues on the PIP-n and the neutral cue on the PIP-r for the 1000ms CD. *“#” denotes group is significantly different than all three neutral groups (blue) based on pair-wise t-tests. Significance based on one-way mixed measures ANOVAs followed by post-hoc pairwise t-tests*.

**Fig 4.**
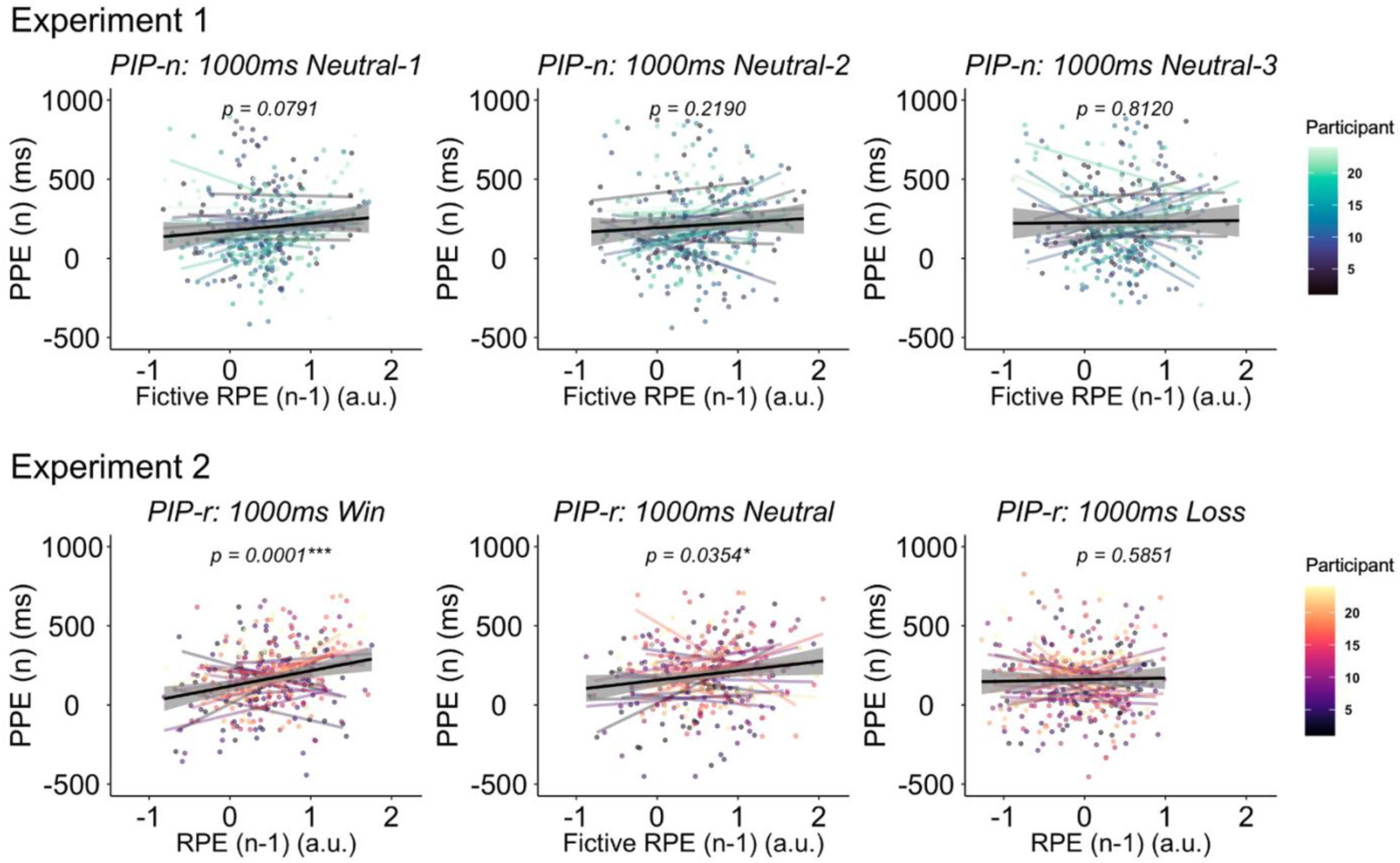
Reward prediction errors (RPEs) on previous trials (n-1) are associated with performance prediction errors (PPEs) on the current trial (n) for positively reinforced cues. Experiment 1. Linear regression models show the results of the association between reward prediction errors of the previous trial (n-1) of the same type and performance prediction errors on the current trail (n) for the 1000ms cues on the PIP-n. **Experiment 2.** for the cues presented on the PIP-r. *Significance based on p < 0.05*, 0.01**, 0.001***. Shading = SEM*.

**Table 1.** Participant demographics and task performance.

|  | **PIP-n (n = 24)** | **PIP-r (n = 24)** |
| --- | --- | --- |
| **Mean age (years)** | 32.92 ± 12.75 | 34.67 ± 14.35 |
| **Gender** | Male: 10<br>Female: 13<br>Other: 1 | Male: 7<br>Female: 16<br>Other: 1 |
| **Handedness** | Right: 17<br>Left: 6<br>Ambidextrous: 1 | Right: 21<br>Left: 0<br>Ambidextrous: 3 |
| **Task time (minutes)** | 22.86 ± 0.57 | 22.13 ± 0.52 |
| **Total points earned** | 107.4 ± 15.34 | 92.45 ± 15.25** |
| **Mean BDI** | 5.750 ± 5.730 | 8.542 ± 7.343 |
| **Mean AMI** | 1.319 ± 0.636 | 1.479 ± 0.832 |
| **Mean QUIP-RS** | 6.916 ± 5.035 | 8.666 ± 5.813 |

*Data expressed as Mean ± SD. PIP = Peak interval procedure. BDI = Beck's Depression Inventory. AMI = Apathy-Motivation Index. QUIP-RS = Questionnaire for Impulsive-Compulsive Disorders in Parkinson's Disease–Rating Scale. Total points equates to the monetary outcome for each task. \*\*Indicates that there was a statistically significant difference in total points between PIP-n and PIP-r participants ( $t(46) = 3.3879$ , $p\text{-value} = 0.0015^{**}$ ). Actual participant compensation was scaled based on performance level and total study time.*

## METHODS

We recruited and consented two groups of 24 participants each from the Winston-Salem, North Carolina region using methods approved by the Wake Forest University School of Medicine IRB (IRB00042265) for a one-hour study visit (**Table 1**). One participant was excluded from the study due to accidental data loss. Sample size was determined by an a priori power analysis. A group-level statistical assessment of the relationship between mean differences in temporal accuracy from a previous study (DiMarco et al. 2023) while anticipating a large (0.8) effect size at power level of 0.8 and p-value of 0.01 for a two-tailed t-test, required recruiting 14-32 participants (per group). These estimates do not use sex or age as factors.

Participants sat approximately 2 feet away from a Dell computer monitor and placed their dominant pointer finger on a button from a hand-held button box. Participants were then instructed on how to play the PIP. The task was designed in Python 3 using the PyGame library. On initialization of the task, nine visual cues were randomly selected from a pool of 60 fractal images sourced from free online stock images. Each cue was assigned an interval of time (3 cues to each category: 1000ms, 3000ms, or 5000ms) (**Fig 1A**, **Fig. 2A**). For the task that included reinforcements, cues were further divided into categories of reinforcement (win, loss, or nothing) (**Fig 2A**). The maximum reinforcement amount was scaled to $3 on all trials on the PIP-r. The reinforcement was calculated using a linear decay of maximum reinforcement in relation to the error in the reproduction time (for example, a reproduction of 950ms for a 1000ms win trial would equate to a reinforcement of +$0.95 whereas a reproduction of 950ms for a 1000ms loss trial will equate to a reinforcement of -$0.05). Cues were randomly presented by placing each cue into an array 16 times and shuffling to determine the sequence of trials for a total of 144 trials.

Stages of the task included: presentation, prompt, reproduction, and reinforcement. During the presentation stage, participants observed the presented cue for the assigned duration (1000ms, 3000ms, or 5000ms) (**Fig 1A, 2A**). Following a black 500ms screen, a ‘ready’ prompt was displayed for 1000ms. After another 500ms black screen, the cue reappeared. Participants were instructed to use the hand-held button box to reproduce the duration of time by pressing the button when the cued duration elapsed. The cue disappeared when the button was pressed or after twice the cued duration passed. Immediately after, a monetary reinforcement or a hairpin cross was displayed based on the reinforcement type. After reinforcement, or if the cue type was neutral, the trial was complete. The cue disappeared, and a black screen was shown for an interval of time randomly drawn from a Poisson distribution with lambda equal to 1500ms. On average, experiments lasted ∼23 minutes per participant (**Table 1**).

Following completion of the task, all participants completed several demographic and clinical questionnaires to better understand and control for the potential individual differences in task performance that have been known to affect timing and reinforcement learning behaviors. These clinical questionnaires included measures of depression (Beck’s Depression Inventory-II), apathy (Apathy-Motivation Index), and impulsivity (Questionnaire for Impulsive and Compulsive Behaviors). At the end of the study visit, participants were compensated based on the study duration ($5 per 15 minutes), with a bonus based on task performance (ranging from $20 - $60 additional) for maximum possible earnings of $80. Analysis was completed using R Studio.

### Experiment 1

To characterize interval timing behavior in response to predictable cues in the absence of reinforcement, we created the PIP-neutral (PIP-n), which presented participants with nine different cues, three of each were predictive of 1000ms, 3000ms, and 5000ms criterion durations, respectively (**Fig 1A**). We recruited 24 participants to complete this task and measured temporal learning using learning curves (**Fig 1B)**. Learning curves were calculated for each participant based on absolute value of the temporal error for each cue type at each appearance, using Eq (1):

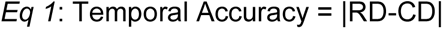

Where RD is the participant’s reproduced duration and CD is the criterion duration. Mean absolute value of the error at each appearance was then fit with an exponential function to determine the rate of temporal learning overtime, using Eq (2):

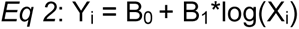

Where Y is absolute error for each cue (i). X is each appearance of the cue (i). B_0_ represents the intercept and measures the value where the line crosses the y-axis. B_1_ is the slope of the curve and measures the rate of change in error or the learning rate.

### Experiment 2

We modified the PIP to deliver reinforcements on 2/3rds of all trials immediately following the participant’s button press, in a task called the PIP-reinforced (PIP-r; **Fig 2A**). During the PIP-r, a reinforcement was a reward (monetary gain) or punishment (monetary loss) immediately following the reproduced duration. The monetary reinforcement structure was the amount of money a participant gained or lost (scaled to ±$3) on a trial and was calculated in a linear fashion from the temporal accuracy of the participant on that trial. Temporal accuracy is calculated using Eq (1). Where RD is the participant’s reproduced duration and CD is the criterion duration (1000ms, 3000ms, or 5000ms). We recruited 24 naive participants to complete the PIP-r and measured their timing behavior using learning curves (**Fig 2B**).

### Reward Prediction Errors

We calculated reward prediction errors as the difference between what participants might have expected to earn on a given cue and what they actually received (**Fig 4**). Reward prediction error (RPE) was calculated using Eq (3):

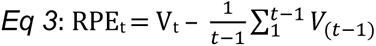

Where t is the current trial with the same cue type (criterion duration x reinforcement type) and V is the monetary value of that trial. Performance prediction errors (PPE) were calculated based on the difference between the expected reproduced duration on a given cue and how participants actually responded to the cue using Eq (4):

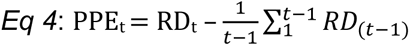

Where t is the current trial with the same cue type (criterion duration x reinforcement type) and RD is the reproduced duration on that trial.

## RESULTS

### Temporal learning without reinforcement

To characterize interval timing behavior in response to predictable cues in the absence of reinforcement we designed PIP-neutral (PIP-n), which presented participants with nine different cues, three of each were predictive of 1000ms, 3000ms, and 5000ms, respectively (**Fig 1A**). Note, the triplicate design is intended to control for the number of cues presented in experiment 2 (PIP-r, described below). We recruited 24 participants to complete this task and estimated temporal learning using fitted learning curves. Learning curves were fit for each participant based on absolute error of reproduced duration (please refer to methods, *Eq. 1*, for calculation of absolute error). Mean absolute error at each appearance was then fit with an exponential function see (methods, *Eq. 2*) to determine the rate of change in error overtime (i.e., temporal learning). The slope of the curve was used to estimate the rate of change in error over repeated trials.

Learning curves for each cued interval on the PIP-n showed significant temporal learning for all criterion durations (**Fig 1B**). Specifically, participants showed significantly lower absolute error at the last appearance (appearance 16) when compared to the first appearance (appearance one) for all cue types (**Fig 1B**; two-way mixed ANOVA; F(1,46) = 10.910; p = 2.00e-03**; generalized eta squared (ges) = 0.081, indicates medium effect size). Mean intercepts of learning curves for each criterion duration also show larger absolute error for longer criterion durations, demonstrating the effects of the scalar property of timing (Rammsayer, & Troche, 2014) (**Fig 1C**; one-way mixed ANOVA; F(2,69) = 12.589; p = 2.18e-05***; ges = 0.267, indicates large effect size). As expected, we did not find significant differences in learning rates or intercepts between neutral cues on the PIP-n within the same criterion duration tested (**Fig S1**). Also, in accordance with other reproduction tasks (Rammsayer, et al., 2015), on average, we observed a general underestimation of all intervals tested (**Fig S1**).

### Increased initial error, but faster temporal learning with positive and negative reinforcement

To test the hypothesis that positive and negative reinforcement of temporal reproduction would increase temporal learning we designed and executed PIP-reinforced (PIP-r, **Fig 2A**). During PIP-r, a reinforcement was paired with a specific temporal cue presented for 1000, 3000 or 5000ms. Reinforcers were a reward (monetary gain) or punishment (monetary loss) delivered immediately following the participant’s reproduced duration. The amount of monetary reinforcement was calculated linearly from the absolute value of the temporal error of the participant on that trial (**Fig 2B**; *Eq 1*; see methods for a description of reinforcement dispersion) scaled to a maximum of ±$3. Three cues indicated a reward, one for each the 1000ms, 3000ms, and 5000ms criterion durations. Another three cues indicated a punishment, one for each the 1000ms, 3000ms, and 5000ms criterion durations. Lastly, three additional cues indicated no reinforcement. We recruited 24 naive participants to complete the PIP-r and estimated their interval timing behavior using fitted learning curves (**Fig 2C**) as described for experiment 1 (PIP-n).

Consistent with behavior on the PIP-n, participants show significantly lower error at the last appearance (appearance 16) when compared to the first appearance for all cues on the PIP-r (**Fig 2C**; two-way mixed ANOVA; F(1,46) = 11.756; p = 0.001**; ges = 0.090). We hypothesized that there would be increased rates of temporal learning in response to the reinforcement schedule on the PIP-r, compared to the lack of reward schedule on the PIP-n. We compared the slope and intercepts of the learning curves from the PIP-n versus PIP-r (**Fig 3 and S3**). We found significantly higher intercepts (**Fig 3A**; mixed ANOVA; F(1,94) = 20.096; p = 2.08e-05***; ges = 0.176), but significantly steeper slopes (**Fig 3B**; mixed ANOVA; F(1,94) = 9.421; p = 0.003**; ges = 0.091) for the neutral cue on PIP-r when compared to the neutral cues on the PIP-n for the 1000ms duration.

Notably, there were no significant differences between the neutral cues on the PIP-n and PIP-r for the 3000ms (**Fig S3B;** mixed ANOVA; F(1,94) = 0.359; p = 0.551; ges = 0.004; **Fig S3E**; mixed ANOVA; F(1,94) = 1.972; p = 0.164; ges = 0.021) and 5000ms (**Fig S3C;** F(1,94) = 1.349; p = 0.248; ges = 0.014; **Fig S3F**; mixed ANOVA; F(1,94) = 3.656; p = 0.059; ges = 0.037) durations. Thus, participants exhibited a specific increase in learning rates in response to reinforced interval timing tasks (PIP-r) for the 1000ms criterion duration but not for the 3000 or 5000ms ms durations.

### Reward Prediction Errors (RPEs) for wins, but not losses, reinforce performance errors

Based on our observation that introducing a reinforcement schedule can increase the initial error (higher intercept of learning curve) but also accelerates temporal learning (increased slope of learning curve) for neutral 1000ms trials and the extant literature, we hypothesized that reinforcement learning mechanisms involving reward prediction errors to actual outcomes and changes in expectations about rewards and punishments drive accelerated learning. Therefore, we calculated RPEs based on the expected value (EV) value of a particular cue (i.e., estimated as their average earnings up to that point on the task for that cue type) and what was actually received on that trial (i.e., the monetary reward or punishment based on accuracy on that cue, see *Eq. 3* in methods). We then fit linear regression models to determine if RPEs on the previous trial are associated with performance prediction errors on the current trial. Performance prediction errors were calculated based on the difference between the expected reproduced duration on a given cue and the actual reproduced duration on the cue (*Eq 4*; described in methods).

From the comparison of RPEs on the previous trial with performance prediction errors on the current trial, we found a significant positive correlation between RPEs on the previous trial and performance prediction errors on the current trial for positively reinforced cues on the PIP-r (**Fig 4, Experiment 2**; linear regression model; F(1,343) = 14.79; p = 0.0001***; R^2^ = 0.0386, indicates large effect size). Interestingly, however, we did not see a relationship between RPEs and PPEs on negatively reinforced cues on the PIP-r (**Fig 4, Experiment 2;** linear regression model; F(1,360) = 0.2985; p = 0.5851; R^2^ = -0.002).

We also calculated ‘fictive-RPEs’ for neutral cues based on what participants *would have accrued if the cues had been reinforced*. We compared previous trial ‘fictive-RPEs’ on the with actual PPEs on the current trial, and we found a significant association for the neutral cues on the PIP-r (**Fig 4, Experiment 2**; linear regression model; F(1,350) = 4.461; p = 0.0354*; R^2^ = 0.0097).

In experiment 1, no reinforcers were provided, yet temporal learning occurred. To determine if the effect of RPE on performance in experiment 2 was specific to reinforcement and not an artifact of general performance improvements we again estimated ‘fictive-RPEs’ based on what participants would have earned if the cues had reinforced and compared the relationship between fictive-RPEs and performance prediction errors during PIP-n. In PIP-r, the RPE is strongly associated with decreases in performance error over time and is a signal that participants observe. On the other hand, the ‘fictive-RPE’ is not observed by participants and should only be associated with the PPE in PIP-n if the effect observed in PIP-r was driven by general performance improvement and not an explicit effect on the actual rewards accrued. We observed no significant associations between fictive-RPEs and performance prediction errors for any cues in PIP-n (**Fig 4, Experiment 1**; linear regression models; Neutral-1: F(1,335) = 3.103; p = 0.07906; R^2^_adj_ = 0.0062; Neutral-2: F(1,343) = 1.513; p = 0.2195; R^2^ = 0.0015; Neutral-3: F(1,339) = 0.0567; p = 0.8119; R^2^ = -0.0028).

## DISCUSSION

The current study investigated temporal learning and interval timing behavior in the presence and absence of reinforcement. Over two experiments, a total of 48 individuals performed a version of a peak interval procedure (PIP) for intervals of 1000ms, 3000ms, and 5000ms criterion durations (**Fig 1 and Fig 2**). The expectation of positive or negative reinforcement was associated with significantly higher initial error (**Fig 3A**), but also was associated with accelerated temporal learning (**Fig 3B**). These effects were specific for 1000ms intervals and were not observed for 3000ms and 5000ms. Notably, a closer examination of the role of a potent reinforcement signal, the reward prediction error (RPE), revealed that trial-by-trial errors, and not performance improvements, were reinforced by ‘actual win’ associated RPEs but not ‘actual loss’ associated RPEs. The effect of ‘actual win’ related RPEs appeared to generalize to the neutral cues (or non-reinforced cues in PIP-r). In model-free temporal difference reinforcement learning theory, cues that acquire the expectation of wins (or losses) are sufficient to drive dopaminergic reward prediction errors. Our results are consistent with the hypothesis that phasic dopamine responses drive expectations associated with cues on the PIP-r, but also to actual wins on the PIP-r driving paradoxical performance changes over the course of the PIP-r. It is possible that early, uncertain expectations increase the salience of cues and drive an initial increase in error in the reproduction of durations. Over repeated trials, temporal learning naturally occurs to overcome this error, but appears to be interfered with by RPEs that reinforce ‘bad’ behavior (**Fig 4**, Experiment 2). Notably, the fastest learning in PIP-r occurs for the non-reinforced neutral cues (**Fig 3B**) where the expectation of an explicit reward (monetary gain) or punishment (monetary loss) is absent.

To understand the effects of reinforcers on interval timing behaviors during an instrumental conditioning task, we first measured interval timing behavior on a PIP in the absence of reinforcement (**Fig 1A**). We showed that participants exhibited temporal learning of 1000ms, 3000ms, and 5000ms intervals of time, as evidenced by the decreased error at the last appearance of the cue compared to the first appearance (**Fig 1B**). We then sought to characterize how reinforcers would affect this temporal learning, so we designed a second experiment to test temporal learning on a PIP in the presence of positive reinforcers, negative reinforcers, and non-reinforced (neutral) cues (**Fig 2A**). We expected temporal learning rates to increase for positive and negative reinforcers, as participants could adjust their behaviors from monetary feedback. However, *within* PIP-r, we saw no differences in learning rates, reproduced durations, or accuracy between reinforced and non-reinforced cues (**Fig 2B**). We hypothesized that this result may be due to the generalization of behavior from reinforced cues to non-reinforced cues on this task. Therefore, we compared temporal learning rates from the PIP in the absence of reinforcement from our first experiment (**Fig 1**) to temporal learning on the PIP in presence of reinforcement from our second experiment (**Fig 2**). We found that the presence of reinforcement on an interval timing task increased temporal learning rates when compared to a task without reinforcement (**Fig 3**). Interestingly, this increase in temporal learning rate was due to an increase in initial temporal error experienced on the reinforced version of the PIP when compared to the non-reinforced version of the PIP.

As the structure of the PIP used to measure interval timing behavior in this study is similar to instrumental conditioning paradigms, it is possible that temporal learning is being accelerated through RPE-mediated mechanisms. Prior to this study, the role of reinforcers as a teaching signal, specifically for interval timing behavior, was unclear. Therefore, we investigated whether RPEs associated with interval timing behavior could help explain differences in temporal learning and temporal errors generated in the context of different valence of reinforcement. Our results show that RPEs on previous trials are associated with temporal performance prediction errors (PPEs) on the current trial for positively reinforced 1000ms cues (**Fig 4**), which suggests that RPEs are reinforcing the poor performance exhibited on the previous trial. Previous research investigating the role of dopamine in interval timing behaviors have led some to propose that reinforcement learning models could be used to explain the dopamine response to interval timing (Petter, et al., 2018; Mikhael & Gershman 2019). As delaying or emitting a reinforcer has been shown to cause the silencing of the dopamine response at the expected time of reward, it has been hypothesized that the dopamine response may track temporal errors as part of or in addition to errors in reward valuation (Hollerman & Schultz 1998). RPE-driven phasic dopamine signals at the outset of each trial and at the end of each trial may be one possible explanation of this increased temporal error but also increased learning rate in the context of a task with reinforcement.

Our study provides evidence that the presence of reinforcement increases the rate of temporal learning for intervals of time of 1000ms in duration (**Fig S3A**), but not 3000ms or 5000ms durations (**Fig S3B-C**). This result is aligned with other research, as different interval timing behavioral effects have been described when comparing durations of less than and greater than approximately 1200ms, with individual differences in the breakpoint (Artieda, et al., 1992; Koch, et al., 2008). It has been hypothesized that different dopaminergic circuitry may be involved in the perception of durations of time less than and greater than approximately 1200ms (Hinton & Meck, 2004). While dopaminergic action in the striatum is thought to play a role in the perception of time of intervals greater than approximately 350ms, the prefrontal cortex and other “decision-making centers” are believed to be engaged when perceiving intervals greater than about 1200ms, as perceiving longer intervals of time is thought to require working memory and temporal planning processes (Smith & Jonides, 1999; Hinton & Meck, 2004; Meck 2005). Our results provide additional evidence of a behavioral difference between timing intervals 1000ms in duration and intervals of greater than 3000ms in duration, as well as may provide some insight into the potential underlying dopaminergic mechanism for intervals of a 1000ms in duration.

Our results demonstrate a significant effect of learning signals that are shared between instrumental learning paradigms and methods used to investigate interval timing behaviors in animal models. In humans, the peak-interval procedure that we used to study interval timing can be instructed and participant performance can be measured, allowing for initial temporal learning effects to be observed. In the TDRL framework there are two key variables that contribute to the calculation of a reward prediction error. We investigate the impact these signals have on temporal learning by comparing temporal learning behavior when these signals are present (PIP-r) versus absent (PIP-n). In TDRL, cues that predict a future possible reward can generate a reward prediction error in the absence of a concurrent reward. It was recently shown that TD -RPEs can elicit a phasic increase in dopamine levels in human striatum (Sands et al., 2023). Together this suggests that the RPE effects observed here may be due to hypothesized phasic increases in dopamine. Interestingly, Sands et al., also showed that dopamine levels rose to the absence of punishment when a punishment was expected (a punishment prediction error), but fell when the punishment was worse than expected. In the current results, the RPE on expected punishment trials failed to reinforce poor performance, but also did not significantly enhance temporal learning. Interestingly, in the context of rewards and punishments, cues that predict neither (the neutral cues in PIP-r) led to significantly faster temporal learning than neutral cues in PIP-n. One possible explanation would be that the neutral cues in the context of PIP-r, gained an expected value that was non-punishing and therefore surprisingly good (better than expected relative to a possibly punishing trial) even though they were not reinforced positively. We did not model the TDRL-RPE in the current work due to significant challenges to modeling the peak-interval procedures without making significant and currently unresolvable assumptions about how the state-space of the task may be represented in the participants’ brains.

Together, our results suggest a significant role of reinforcement learning mediated dopamine signals in time perception. Others have suggested such a role (Gershman, et al., 2014; Petter, et al., 2018; Mikhael & Gershman, 2019; and Jakob, et al., 2022), but have proposed models that deviate from the simplest and most well-established model for dopaminergic neuron activity and dopamine release (i.e., TDRL representations). Conceptually, while TDRL models reference *time* in its name and in its indexing of variables, *time* is not per se necessary for calculating reward expectation errors or updates in expected values. The framework classically uses *time* simply as an intuitive indexing variable to signify different sequences of states the agent or environment may evolve through. As such, the classic TDRL framework only requires that the agent keep record of what state it is in and what state is likely *next*. A particular interval of time is not necessary to be maintained – *next* could be a few milliseconds, minutes, days, or years. Thus, if the brain or a computerized agent were to simply keep track of a dynamic state-space with the ‘current’ state associated probabilistically with a state-transition array, then actions (like reporting the end of a *time* interval) could be learned through reinforcement learning mechanisms where the representations are not *time* intervals, but instead probabilities that acting in a given state are associated with a given expected positive value. In this manner, *time,* as we perceive it, could be represented without an explicit reference to *time*, but rather a representation that can signify ‘state’ and most likely ‘different state’.

Our suggestion that the time we perceive may be represented formally without reference to time as a variable using the TDRL framework is hypothetical and requires more work to determine and explicate in mathematical form. Additionally, future work should be aimed at understanding whether the expected value and reward prediction error effects observed here in humans is indeed mediated by associated changes in dopamine in human striatum. Nevertheless, we have demonstrated that error signals, previously demonstrated to modulate phasic dopamine levels, can modulate timing behavior and temporal learning in paradoxical ways. Without directly measuring dopamine during these behaviors in the absence of confounding behavioral training, it will be difficult to tease out the role dopamine plays specifically in timing behaviors and temporal learning versus effects that are likely associated with instrumental learning paradigms requires to have non-human animals perform these behaviors. While dopamine, reward prediction errors, and timing behaviors are clearly interconnected, significant work remains to develop a formal model of the time we perceive that does not circularly self-reference the key variable of interest.

## SUPPLEMENTAL FIGURES

**Fig S1.**
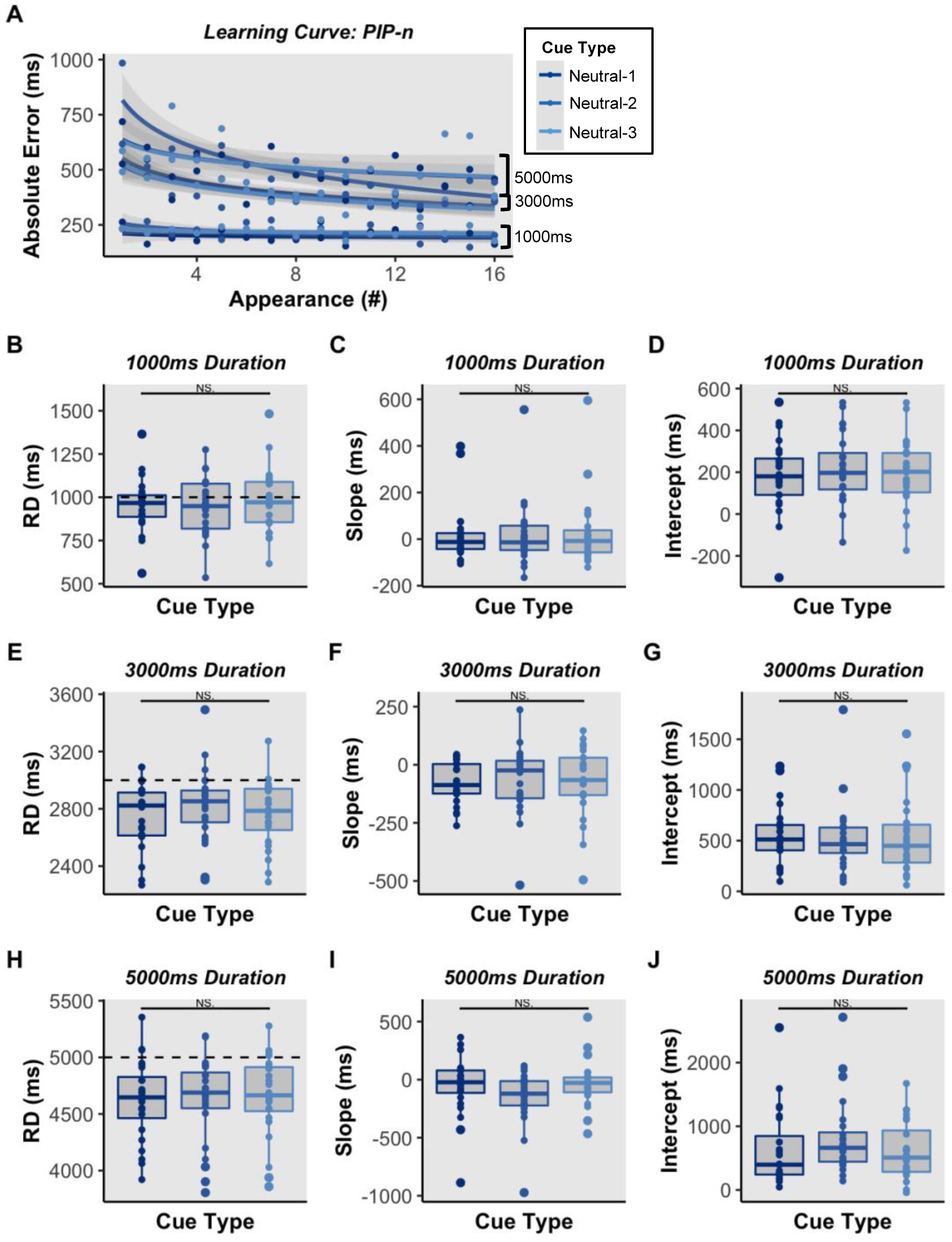
Interval timing is similar across all neutral cues on PIP-n. **A.** Mean learning curves for all neutral cues on PIP-n. **B.** No significant differences between mean reproduced duration (RD) for 1000ms neutral cues. **C.** No significant differences between mean learning rate (slope) for 1000ms neutral cues. **D.** No significant differences between mean intercept for 1000ms neutral cues. **E-G.** For 3000ms duration. **H-J**. For 5000ms duration. *NS. indicates not significant based on one-way mixed ANOVA*.

**Fig S2.**
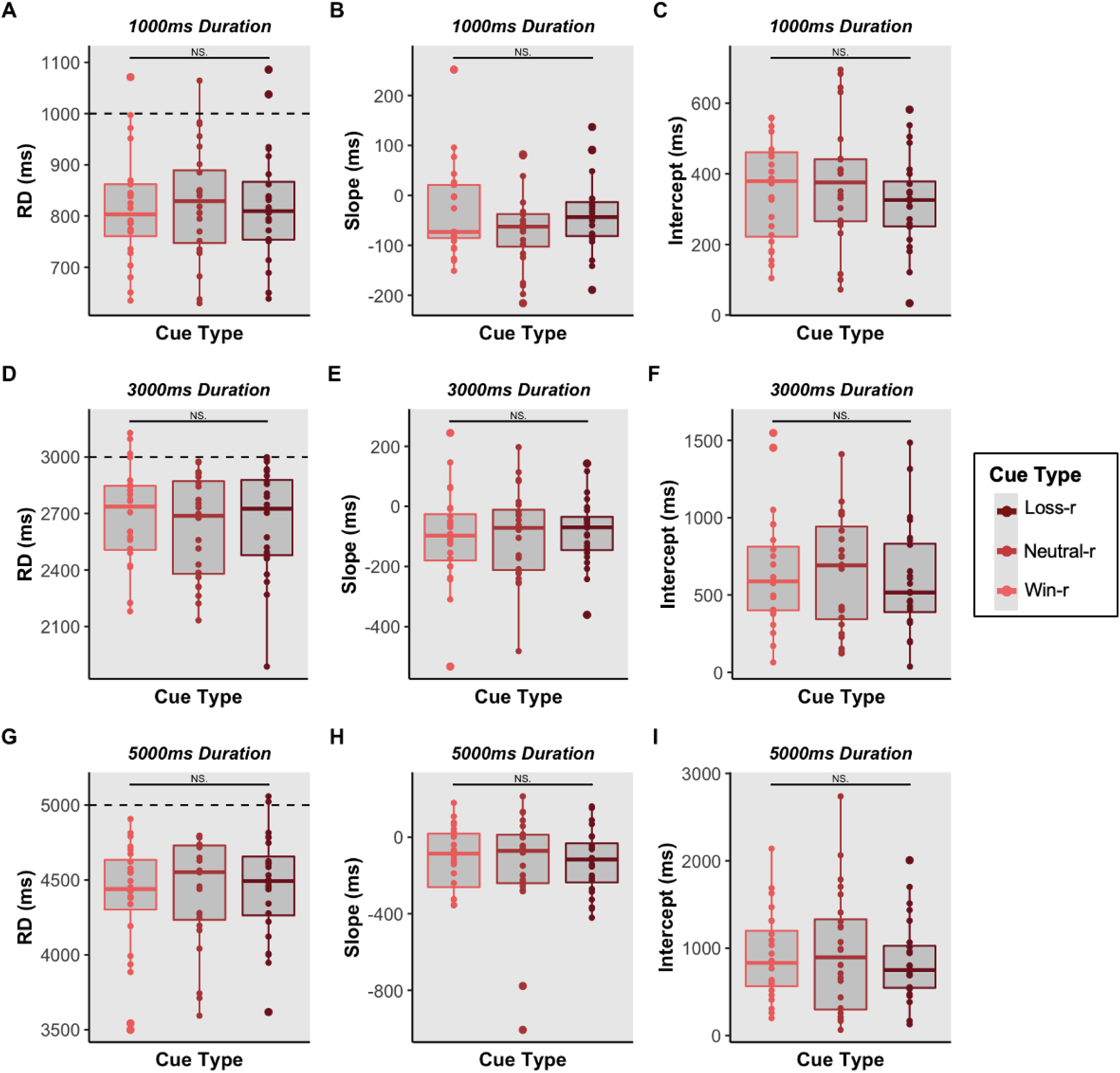
Interval timing is similar across all cues on PIP-r. **A.** No significant differences between mean reproduced duration (RD) for 1000ms cues. **B.** No significant differences between mean learning rate (slope) for 1000ms cues. **C.** No significant differences between mean intercept for 1000ms cues. **D-F.** For 3000ms duration. **G-I**. For 5000ms duration. *NS. indicates not significant based on one-way mixed ANOVA*.

**Fig S3.**
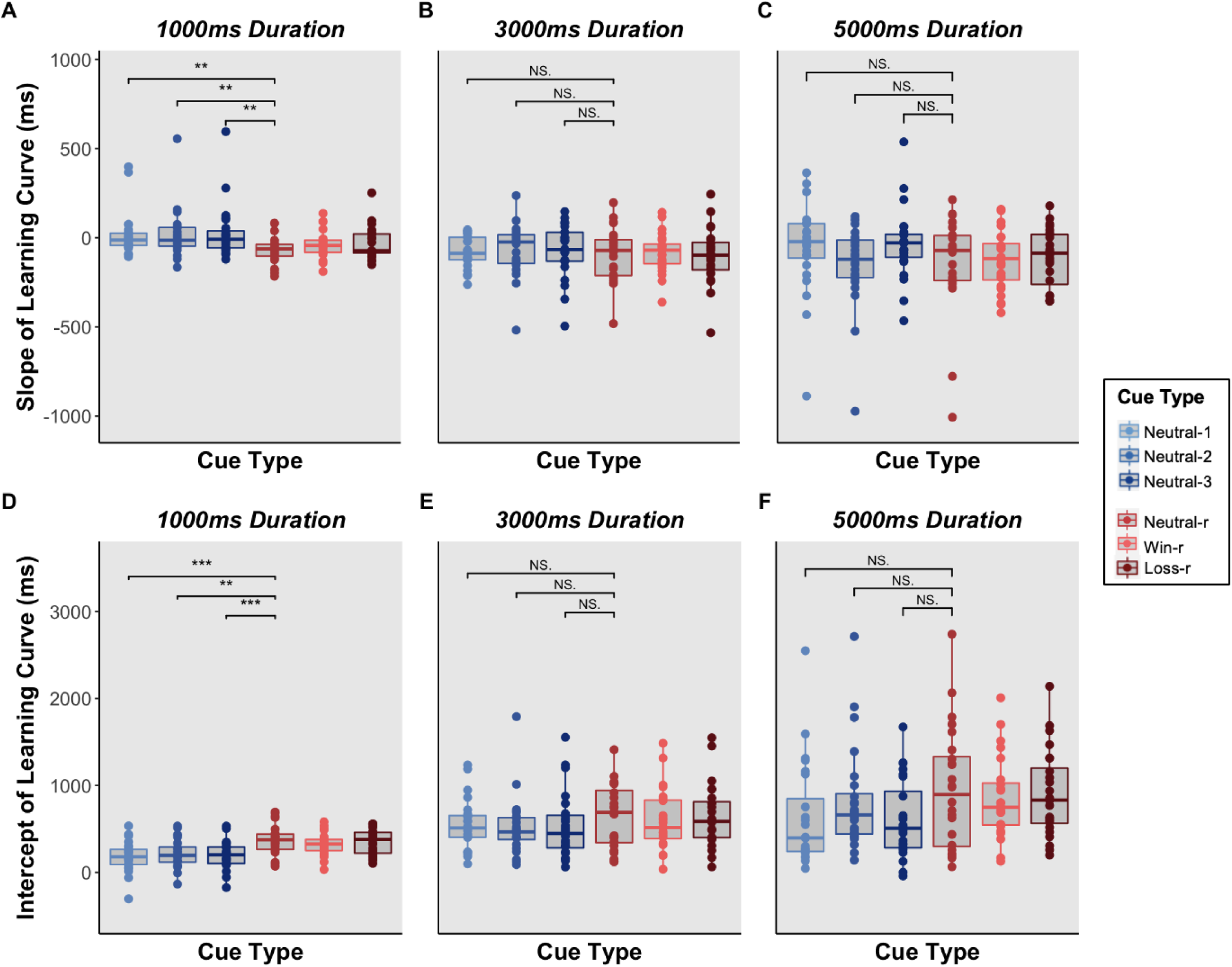
Learning rates differ between reinforced and non-reinforced cues for the 1000ms duration, but not the 3000ms and 5000ms durations. **A.** Comparing the slopes of the learning curves from the neutral cues on the PIP-n (blue) to the neutral cues on the PIP-r (red) for the 1000ms duration. **B.** for the 3000ms duration. **C.** for the 5000ms duration. **D.** Comparing the intercept of the learning curves from the neutral cues on the PIP-n (blue) to the neutral cues on the PIP-r (red) for the 1000ms durations **E.** for the 3000ms duration. **F.** for the 5000ms duration. *Significance based on p < 0.05*, 0.01**, 0.001***. NS. indicates not significant based on one-way two-sample t-tests*.

**Fig S4.**
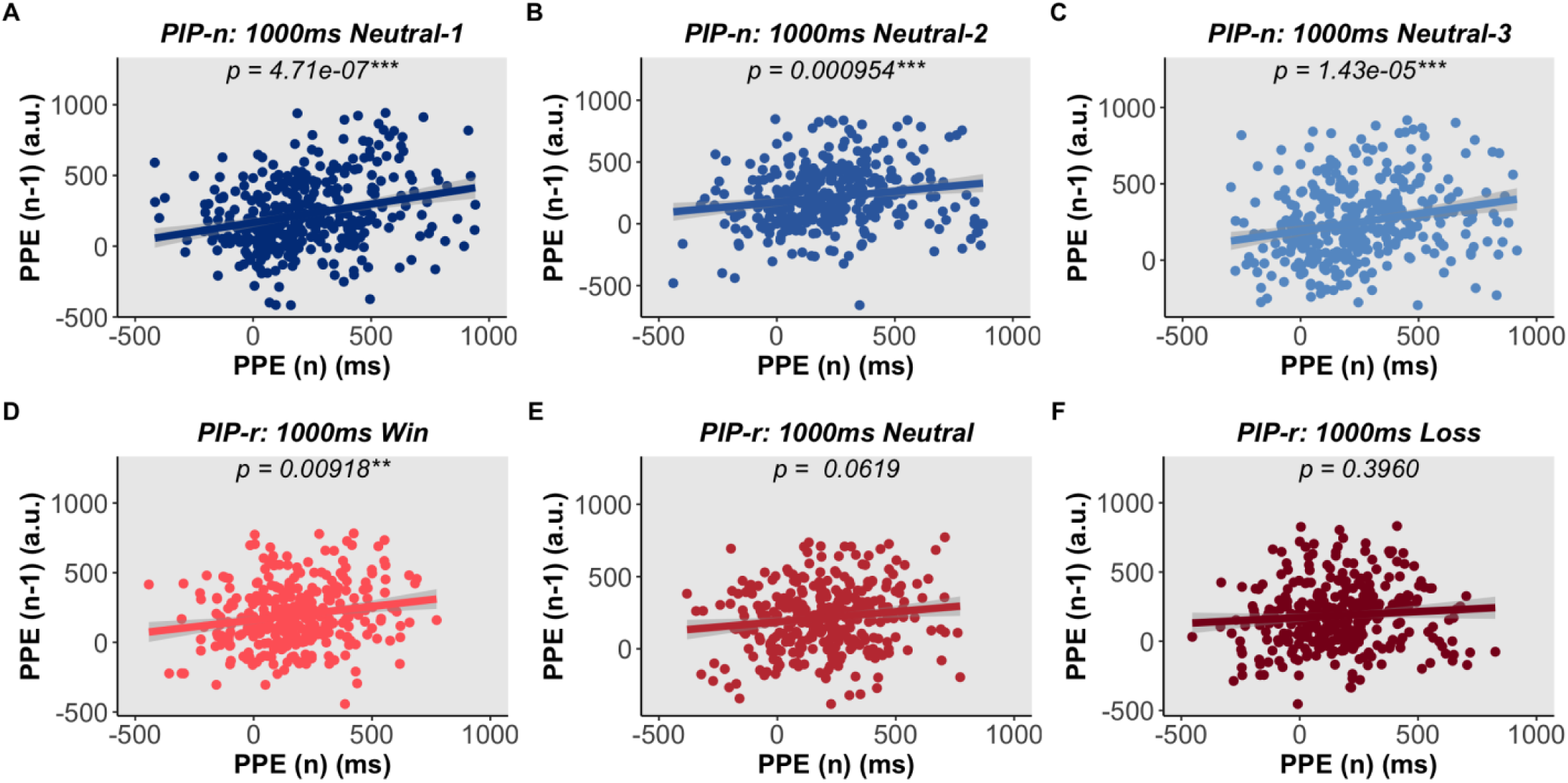
Performance prediction errors on previous trials (n-1) are associated with performance prediction errors on the current trial (n), except for the 1000ms loss cue on the PIP-r. **A.** Linear regression models show the results of the association between performance prediction errors of the previous trial (n-1) of the same type and performance prediction errors on the current trail (n) for the 1000ms neutral-1 cue on the PIP-n. **B-L.** for the other cues presented on the PIP-n and PIP-r. *Significance based on p < 0.05*, 0.01**, 0.001***. Shading = SEM*.

